# Environmental niche forecasts of globally threatened yellow-throated bulbul, Pycnonotus xantholaemus for conservation prospects in the Deccan peninsula, India

**DOI:** 10.1101/633214

**Authors:** Ashish Jha, Karthikeyan Vasudevan

## Abstract

Yellow-throated Bulbul, *Pycnonotus xantholaemus* is an endemic, rare and threatened species with discontinuous distribution across Deccan peninsula. IUCN has enlisted surveys for newer populations as one of the conservation measures. We used maxent algorithm to generate environmental niche model for further surveys. We looked into climatic envelope at occurrence points and contrast it with background climatic envelope. We compared the model for current scenario and future scenario to assess change in extent of predicted niche over time. We used six variables: climatic, topographical and vegetation layers, and a final set of 102 verified presence locations to generate the model. Topographic ruggedness index and precipitation of wettest month (Bio13) were the strong predictors for Yellow-throated Bulbul niche. Model predicts highly discontinuous and small fragments totalling 7% area of peninsular India as suitable niche. Only 10 % of predicted niche falls within India’s protected area network. Loss of habitat due to granite quarrying and anthropogenic pressure will be a bigger threat than climate change.

## Introduction

Yellow-throated Bulbul, *Pycnonotus xantholaemus* (here after YTB) is one of the 93 threatened species of bird in India [1, 2, Fig_1]. YTB is a passerine endemic to scrub forests of the Eastern Ghats, inland hillocks and eastern slope of the southern Western Ghats in the Deccan peninsula. It is speculated to occur in similar habitats in the State of Odisha, but there are no reports to confirm this [3, 4]. It is restricted to scrub forests along the hill slopes and has patchy distribution across its geographical range, with a close association to rocky outcrops [3, 5]. Therefore, quarrying of these granitic rock amounts to habitat destruction for this species. Owing to its discreet population and ongoing loss of habitat, it has been classified as ‘vulnerable’ in IUCN redlist [1]. Currently, there is no conservation action plan or monitoring program in place for this species. The species has been overlooked because of its skulking behaviour [6] and its habitat does not receive the desired levels of protection due to lack of charismatic large mammalian vertebrates [7]. Most of the information on its distribution and natural history comes from opportunistic incidental sightings and available literature on the species is limited to field guides and locality records. Focussed species surveys, estimation of population, lobbying against large-scale granite quarrying and creating awareness have been proposed as conservation measures for this species [1].

**Fig_1.**
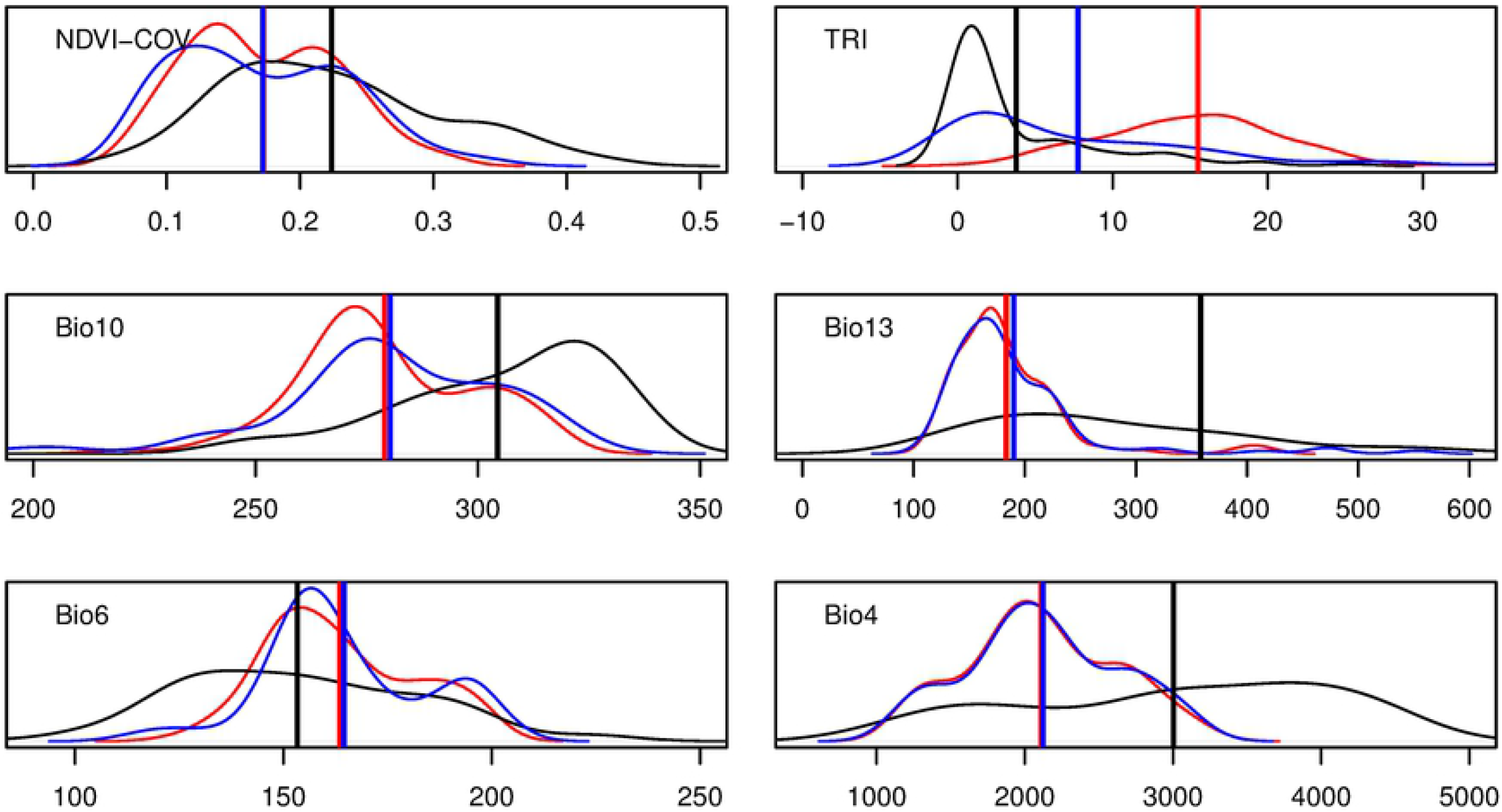
Environmental niche model for Yellow-throated Bulbul. Model generated across peninsular India is shown in grey shading. The lines demarcate the State boundaries. Black dots represent the occurrence records used to generate the model. Enclosed boxes represent: a) Northern Western Ghats, b) Northern Eastern Ghats, c) Southern Western Ghats showing prominent hill ranges, d) Yellow-throated Bulbul, e) Map of India showing the Eastern and Western Ghats, and the Maxent background. The figure was made using ArcGIS 10.3.

Knowledge about the geographical distribution and the preferred habitat of a species is crucial for successful species conservation [8]. For rare species, ecologists have relied upon various Species Distribution Models (SDM) to delimit environmental niche and to form conservation strategies [9–14]. SDM are predictive models which relate presence locations with suits of environmental variables to create an environmental niche map [15]. Some SDM approaches require presence – absence data for model construction while others require presence-only data [13]. Maximum Entropy (MAXENT) method is a presence-only modelling approach that has been widely used to implement SDM [16, 17]. Maxent offers a useful application for determining the environmental niche of rare and elusive species, such as the YTB. Further, ascertaining absence in a location is always tricky and conducting systematic field surveys across a large geographical range is expensive.

In this study, we generated maxent based SDM of YTB in the Peninsular India, and: 1) estimated extent of the environmental niche in the region; 2) measured the difference in environmental niche of the species from the background and 3) forecast climate change impacts on the environmental niche and evaluate anthropogenic threats to YTB habitat.

## Materials and Methods

### Study area

It was in peninsular India bounded within 72°E–88°N and 8°N–22°N. This region comprised biogeographic province 5A (Western Ghats- Malabar Plains), 5B (Western Ghats- Western Ghat Mountains), 6C (Deccan peninsula- Eastern Highlands), 6D (Deccan peninsula- Central Plateau), 6E (Deccan peninsula- Deccan South) [18]. Deccan plateau is hemmed between two mountain ranges- Eastern Ghats and the Western Ghats (see inset e in Fig_1). The Eastern Ghats constitutes a large part of Deccan plateau and it is located on the leeward side of the Western Ghats. It receives an annual rainfall ranging from 20-100 cm [19] in the leeward side of Western Ghats and it typically has dry deciduous and scrub vegetation. The global presence localities of YTB are restricted to the Deccan peninsula.

### Occurrence records

We made field visits to known and other probable YTB locations across four states in Peninsular India during 2015-2018 and recorded YTB in 28 locations (S1_Table.docx). We collected first-hand information on their habitat, behaviour, ecology and recorded habitat features such as, elevation, vegetation, presence of water, rocky outcrops, and anthropogenic pressure. With popularity of bird photography and bird watching, several locations of YTB have been added on online platforms, such as, eBird [20], Flikr, and Oriental Bird Images in recent times. We collated, in all, 202 reported locations from literature, informants, local naturalists, and online platforms. All the reports compiled from secondary sources were screened for reliability. The GPS points of each location were projected using Google Earth™ and the validity of the location was assessed. A location was considered for the study, if: (i) the species was reported by multiple and independent observers; (ii) there was a photograph of the species or there were detailed field observations made by the observer and; (iii) topography of the location matched the description of YTB habitat. In case of any ambiguity, we contacted observers and sought more information about the location, before it was included in the study. Spatial thinning was performed manually in ArcGIS 10.3 to remove duplicate occurrences and reduce the sampling bias. All selected locations were chosen in a manner that they were spaced apart by > 2 km to reduce spatial autocorrelation. A final set of 102 presence points were used for niche modelling (Table_S1.docx).

### Environmental variables

We obtained bioclimatic (WorldClim dataset), topographic (Digital elevation map) and vegetation layers (NDVI layers) at a resolution of 1km^2^ from online sources (Table 1) and the layers were clipped for Peninsular India in ArcGIS 10.3. Based on field observations and knowledge of the biology of the species, we selected layers which were biologically and ecologically relevant. Out of the 19 bioclimatic variables, four were used for final analysis after removing variables that either had Pearson’s r > 0.75, or those with < 2% contribution to the preliminary model. The selected four bioclimatic variables were used for IPPC5 climate projections from global climate models (GCMs - CCSM4). Moderate climate scenario - Representative Concentration Pathway 4.5, was used to project the model for 2050.

**Table 1.** List of predictor variables used in Maxent analysis, with percent contribution and permutation importance in predicted niche model.

| Variable | Source | Percent contribution/<br>Permutation importance |
| --- | --- | --- |
| TRI; Topographic Ruggedness Index (derived from Digital elevation map) | [23] | 59.9/ 58.1 |
| Bio13; Precipitation of Wettest Period (mm) | [21] | 20.5/ 31.9 |
| Bio4; Temperature Seasonality (SD*100) | [21] | 15.4/ 6.7 |
| Bio10; Mean Temperature of Warmest Quarter (°C*10) | [21] | 2.3/ 1.9 |
| NDVI-COV; Coefficient of Variation of NDVI Layers, Jan 2011-Jan 2019 | [22] | 1.1/ 0.4 |
| Bio6; Min Temperature of Coldest month (°C*10) | [21] | 0.9/ 1.1 |

On the basis of field surveys, we observed that YTB is closely associated with presence of water seepage and rock formations. The rock formations constituted granitic gneisses, the charnockite series, the khondalite series and the granites [24]. The seepages in the rock formation are usually perennial with dense vegetation. These habitat patches were mesic in nature, which supported a variety of trees, and shrubs. Some of them were food plants of the YTB for e.g., *Ficus amplissima, F. microcarpa, F. mollis, Erythroxylum monogynum, Premna tormentosa* (personal observation). The mesic patches were < 10 ha, had equitable ambient temperature and contained perennial source of water, in an otherwise hot and arid landscape. We used NDVI data of year 2011-2018 to create a layer of coefficient of variation using raster calculator in ArcGIS 10.3 to capture this important feature of YTB habitat. A low coefficient of variation would indicate that the area experienced equitable abiotic conditions throughout the year. Our preliminary models had a large weightage to elevation and they did not predict the niche in some prominent locations in low elevations. The models instead predicted the niche in high elevation (plateau), where there were no reports of YTB. Therefore, we created a layer of topographic ruggedness index using digital elevation map in ArcMap 10.3, and used it in the final analysis, instead of the elevation layer.

### Modelling procedure and evaluation

Output format was set to logistic, output file type was ASCII and 30% of occurrence points were set aside as the testing dataset. Bootstrap method was used as replicated run type and maximum iteration was set at 5000. All other settings were set as default. We used the average model of fifteen replicates for the subsequent analysis. We used maximum training sensitivity and specificity threshold, to transform maxent’s non-binary distribution model into binary prediction [25].

To visualize how the climatic envelope at occurrence points differs from random points, we generated 102 random points across the background and within 10 km radius of occurrence point and plotted density distribution of the variables at these points and verified occurrence records. To understand the impact of climate change on the environmental niche of YTB, the same set of predictor variables were used to re-run Maxent for current and future scenario. Since, topographic ruggedness index, a topographical feature will not change by 2050 from the present, it was used as such. Vegetation data (NDVI) are unavailable for future projections, hence we re-ran maxent model using five variables for current and 2050 scenario, excluding vegetation layer. Additionally, we re-ran Maxent for both the scenario using vegetation layer assuming no change in vegetation structure in YTB habitats in 2050. The change in environmental niche was calculated in ArcGIS 10.3 using raster calculator by subtracting raster output obtained from Maxent algorithm. The resulting raster was reclassified into ‘increment’, ‘no change’ and ‘decrease’ in environmental niche.

The area under the curve (AUC) of the receiver operating characteristic (ROC) plot [16] and the true skill statistics (TSS) [26] were utilized to assess the model’s explanatory power. AUC measures the model’s ability to distinguish between the presence records and the random background points. AUC values ranged from 0.5 (not different from a randomly selected predictive distribution) to 1 (with perfect predictive ability). The TSS is defined as one minus sum of sensitivity and specificity, and it ranges from −1 to 1. A value of one indicates perfect agreement; a value of zero or lesser indicates a performance no better than random; a value of minus one indicates perfect disagreement. We also tested significance of the model using the null model approach [27]. We generated 99 sets of 102 random points without replacement across the background and the minimum convex polygon. We ran maxent to calculate AUC for both the null distributions and compared the 95th percentile of AUC value of the null-distributions with average AUC value of the model. Model was considered significantly better than the null, if the AUC value exceeded the 95th percentile of the null AUC distributions.

## Results

### Predicted Environmental Niche of YTB

The model scored mean TSS 0.909±SD 0.023 and mean AUC 0.975±SD 0.0063 with species’ model AUC being significantly better than random (Fig_S1). Maximum sensitivity plus specificity threshold (0.0994) was used to convert maxent output to binary layer. Maxent predicted an area of 1,03,790 km^2^ as the suitable environmental niche. This area is distributed in a disjunct manner, with 80% area in fragments of < 5km^2^. In addition, to localities in and around known occurrence points, maxent predicts suitable niche outside the recorded limits in following three locations:(a) 3065 km^2^ in the Northern Western Ghats; (b) 19,534 km^2^ in the Northern Eastern Ghats and; (c) 5821 km^2^in the Southern Western Ghats (Fig_1). Predicted model in Ascii format and supporting files have been provided (S_Maxent_output.zip).

### Variables determining the environmental niche of YTB

Topographic ruggedness index appears to be most important predictor followed by Precipitation of Wettest Period (Bio13) based on jackknife test and permutation importance of the environmental variables (Table 1). NDVI-coefficient of variation layers affects maxent prediction as expected. Response curve shows that with increase in values of coefficient of variation layers, logistic output decreases (Fig_S2). The distribution and mean value of variables at the occurrence locations differ from those of the background. The locations where the bird occurs are topographically more rugged than the background area. Precipitation of wettest period (Bio13), temperature seasonality (Bio4), mean temperature of warmest quarter (Bio10), and NDVI-coefficient of variation is lower at locations where the bird occurs than the background. The minimum temperature of coldest month (Bio6) is higher in locations where it occurred, than in the background locations (Fig_2).

**Fig_2.**
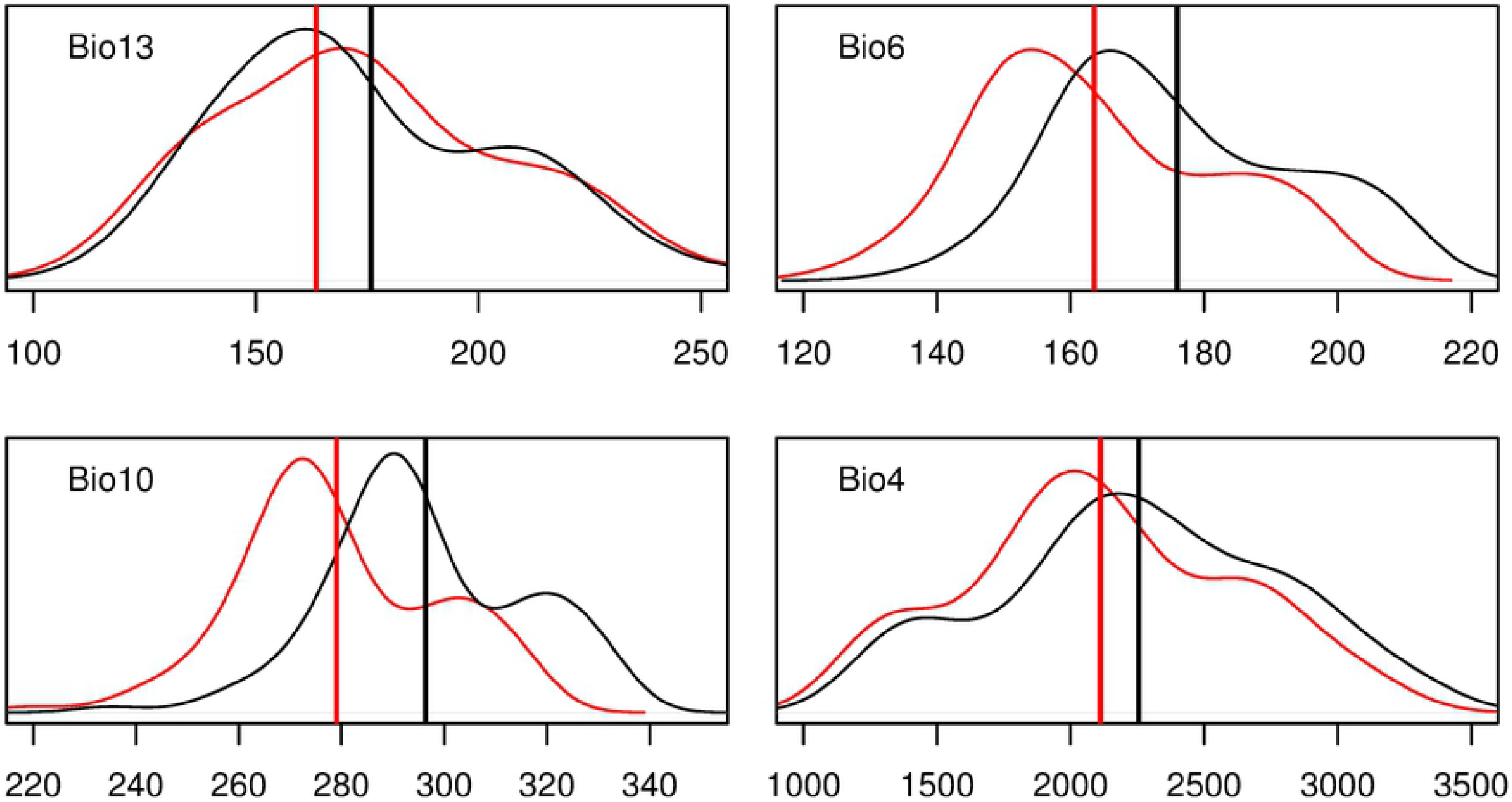
Density distribution of bioclimatic variables at verified occurrence points and background. Density plots generated at 102 verified YTB occurrence points (red), 102 random points within 10 km radii of verified occurrences (blue) and 102 random points across the background (black). Raster values at the points were extracted in ArcGIS 10.3 and imported in R 3.1.2. Straight line represents the mean value of the distribution. The x and y axes represent range and density respectively. Refer table 1 for the description of variables.

### Predicted environmental niche of YTB due to climate change

The model scored mean TSS 0.888±SD 0.023 and mean AUC 0.973±SD 0.0054 for 2050. Maxent predicts marginal changein extent of suitable niche of YTB in 2050. Change in extent of suitable niche was comparable for: a) both current and 2050 scenario with five predictor variables, excluding vegetation data and b) both current and 2050 scenario with six predictor variables, including vegetation data. Predicted shift in mean values of bioclimatic variables in 2050 is within the distribution range of bioclimatic variables at current scenario (Fig_3). Few regions, particularly Southern Western Ghats show marginal increase in extent of environmental niche in 2050 towards elevation close to 1000 m.

**Fig_3.**
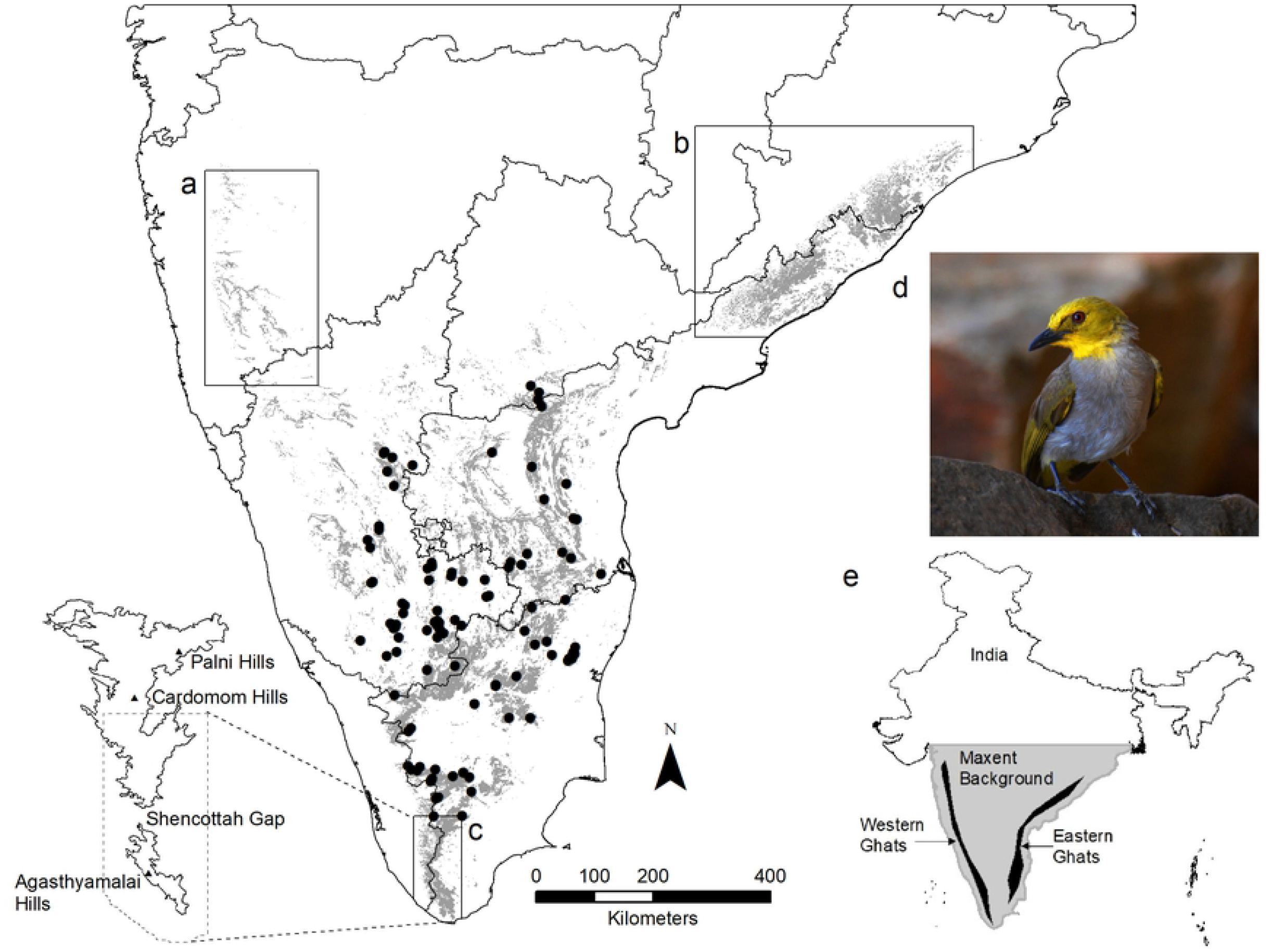
Density distribution of bioclimatic variables for current and 2050 scenario. Density plot at 102 verified YTB occurrence points for current scenario (red) compared with 2050 scenario (black). Raster values at the points were extracted in ArcGIS 10.3 and imported in R 3.1.2. Straight line represents the mean value of the distribution. The x and y axes represent range and density respectively. Refer table 1 for the description of variables.

## Discussion

### Predicted Environmental Niche of YTB

Model predicts suitable niche along the Eastern Ghats, inland hillocks and dry eastern slopes of the Western Ghats. Only 10 % of predicted niche falls within the protected area network (Table_S1.docx). The prediction of the environmental niche in the northern Eastern Ghats (Fig_1b) warrants systematic surveys in the region. There is only one record of YTB from this location [28]. The model also predicts suitable environmental niche at Southern Western Ghats (Agathyamalai Hills). The bird is recorded from adjoining hills, Cardamom Hills and Palani Hills, which lies north of Shencottah gap while Agasthyamalai hills lies south of the gap (Fig_1c). Suitable habitats were also predicted along the northern Western Ghats however nearest known occurrence is 300 km east in Bellary, Karnataka. The bird has been reported from eastern slopes of Western Ghats. It should be noted that eBird has few hotspot records from above mentioned localities, and other locations are not frequently visited by birders. Given the rarity of the species, its skulking habit, and the lack of birding effort in these locations, it is possible that these locations have hitherto, populations of YTB that have not been recorded. A list of locations for future surveys for the species (Table_S2.docx) would inform birders in the region.

### Variables determining the environmental niche of YTB

Among the selected variables, topography differed the most in locations where YTB was found in comparison with random locations within 10 km radii of the occurrence locations (Fig_2). Topography of the landscape would further impact other bioclimatic features such as moisture retention and temperature regimes. This suggests that though the species has a large geographic range, its distribution is influenced by abiotic conditions, and topography associated features. Since the environmental niche of YTB is also a refuge against extremely high temperatures, hosts fruiting trees, and water source, it might attract several other species to use this location in the landscape.

### Impact of climate change and habitat loss on YTB

Projection of SDM in future climate change scenario shows only a marginal change in extent of suitable niche. This could be due to the wide latitudinal range (9°N – 16° N) and the variation is habitat characteristics and precipitation in the region. The Model predicts a marginal rise in extent of suitable habitat in Western Ghats. At present, the species is reported from Western Ghats up to an elevation of 900 m, and beyond this elevation, the vegetation transitions into the evergreen type. Under climate change scenario, this region is expected to get warmer [29], thereby, allowing the species to occupy elevations beyond its current upper elevation limit. Niche shifts due to climate change based on bioclimatic projections are insufficient to infer on the area of occurrence in 2050. Factors such as, physiology, life history traits, and anthropogenic pressure would ultimately determine the future geographic range of a species [30]. Since YTB is a behaviourally timid species [31, 3], climate change driven range shifts might expose it to competition from sympatric red-vented bulbul *Pycnonotus cafer*, and white-browed bulbul *P.luteolus*.

Surging population in India is expected grow till 2050 [32]. The Eastern Ghats alone in the Deccan Peninsula has undergone 40% reduction in forest cover between 1920–2015 [33] and the human population is predicted increase in the region [34]. Twelve out of 27 sites with YTB occurrence recorded by us had religious tourism. If this activity is left uncontrolled, it will threaten YTB habitats. The demand for cultivable land and mining of natural resources such as granite and construction aggregate from rocky outcrops are likely to grow. Construction aggregate is an important construction material and it is also mined from rocky outcrops. The states those encompass the geographic range of the species also holds India’s 25% granite reserves in the rocky outcrops. Therefore, quarrying for granite is expected to expand in the region [35]. Demand for construction aggregate is also expected to soar beyond 2 billion tonnes by 2020 [36]. The environmental niche of YTB is undoubtedly threatened due to anthropogenic pressure from quarrying in the entire Deccan Peninsula.

### Conservation

YTB were sighted at few locations with active human presence (S3_Table.docx). It appears that low intensity human activity does not impact the species, as long as scrub forests are retained along the hill slopes and rocky outcrops. Therefore, conservation of YTB is possible alongside human use of the Deccan Peninsula.

Though YTB has a large area of occurrence in the Deccan peninsula, it inhabits very small habitat patches, and it is found in densities lesser than the other sympatric bulbuls [5]. Such small and scattered populations make them vulnerable to stochastic extinctions [37]. Mapping YTB populations at regional level, estimation of population size, the study of movement and genetic structure in the population will generate vital information for its conservation. Given the close association of the species with large, exposed basaltic outcrops in the Deccan peninsula, it can serve as a flagship species for this biogeographic zone to consolidate the remnant habitats and hasten reclamation of abandoned quarry sites within geographic distribution of the species.

## Acknowledgements

We thank Director CSIR-CCMB for providing funds for this study. We acknowledge H.S. Prayag, Karnataka Agriculture University, Bengaluru, for assistance during fieldwork at Karnataka. We extend our acknowledgement to the state forest departments of Telangana, Andhra Pradesh, Karnataka and Tamil Nadu for having given permissions to conduct field studies. We acknowledge Colorado State University, USA, National Centre for Biological Sciences, Bengaluru and Indian Statistical Institute, Kolkata for training AJ through workshops in species distribution modelling.

## Author Contributions

AJ and KV conceived and designed the experiments, and wrote the paper. KV supported the study through CSIR-CCMB funds. AJ conducted field work, collected the data, and analysed the data.

## Supporting information captions

**Fig_S1. Histogram of AUC of null distributions.** Plot shows AUC distribution of 99 independent maxent models run using 102 random records obtained from background (grey bars) and from Minimum convex polygon around the verified occurrences (black bars). Arrow shows mean AUC value of maxent model run using verified occurrences.

**Fig_S2. Response curves of variables used for modelling.** Refer Table 1 for the description of variables.

**Table_S1. State wise locations of Yellow-throated Bulbul used for the maxent modelling.** Note: Protection status enlists areas designated under Wildlife Protection Act, 1972 as Wildlife Sanctuary (WLS)/ National Park (NP) or managed by Archaeological survey of India (ASI) or state Forest Department.

**Table_S2. Tentative locations for survey of Yellow-throated Bulbul.** *Ghat* roads, Hill Shrines, and Waterfalls within 1000 m elevation with scrub forests in below mentioned locations are most promising places to survey for the species.

**Table_S3: Few prominent YTB locations with high anthropogenic activity.** Details of locations given in S1_Table.

## References

1. BirdLife International. Pycnonotus xantholaemus. The IUCN Red List of Threatened Species 2016: e.T22712719A94345114. Downloaded on 15 February 2019. Available from: http://dx.doi.org/10.2305/IUCN.UK.2016-3.RLTS.T22712719A94345114.en.

2. Praveen J, Jayapal R, Pittie A. Threatened birds of India (v2.1). 2019 January 15 [cited 11 March 2019]. Available from: http://www.indianbirds.in/india/.

3. Ali S, Ripley SD. Handbook of the birds of India and Pakistan. Compact ed. New Delhi: Oxford University Press; Vol 6: 93–96, 1987.

4. Subramanya S. Does the Yellow-throated Bulbul *Pycnonotus xantholaemus* occur in Orissa? Newsletter for Ornithologists. 2004; 1: 39–40.

5. Subramanya S, Prasad JN, Karthikeyan S. Status, habitat, habits and conservation of Yellow-throated Bulbul *Pycnonotus xantholaemus* (Jerdon) in South India. Journal of Bombay Natural History Society. 2006; 103: 215–226.

6. Ali S. The birds of Mysore. Journal of Bombay Natural History Society. 1942; 43:318–41.

7. Rahmani, A. Editorial: Threatened birds of India: need for immediate conservation action. Journal of Bombay Natural History Society. 2010; 106: 1–3.

8. Margules CR, Pressey RL. Systematic conservation planning. Nature. 2000; 405: 243–253.

9. Peterson, AT, Robins CR. Using ecological-niche modelling to predict barred owl invasions with implications for spotted owl conservation. Conservation Biology. 2003; 17: 1161–1165.

10. Raxworthy CJ, Martínez-Meyer E, Horning N, Nussbaum RA, Schneider GE, Ortega-Huerta MA, Peterson AT. Predicting distributions of known and unknown reptile species in Madagascar. Nature. 2003; 426:837–841.

11. Sánchez-Cordero V, Cirelli V, Munguía M, Sarkar S. Place prioritization for biodiversity content using species ecological niche modelling. Biodiversity Informatics. 2005; 2, 11–23.

12. Hernandez PA, Graham CH, Master LL, Albert DL. The effect of sample size and species characteristics on performance of different species distribution modelling methods. Ecography. 2006; 29: 773–785.

13. Tsoar A, Allouche O, Steinitz O, Rotem D, Kadmon R. A comparative evaluation of presence-only methods for modelling species distribution. Diversity and Distributions. 2007; 13: 397–405.

14. Kumara HN, Ullah MI, Kumar S. Mapping potential distribution of slender Loris subspecies in peninsular India. Endangered Species Research. 2009; 7:29–38.

15. Guisan A, Zimmermann NE. Predictive habitat distribution models in ecology. Ecological Modelling. 2000; 135: 147–186.

16. Phillips SJ, Dudík M, Schapire RE. A maximum entropy approach to species distribution modeling. – In: Brodley, C. E. (ed.), Machine learning. Proceedings of the twenty-first international conference on machine learning, Banff, Canada, ACM Press; 2004. pp. 83.

17. Elith J, Phillips SJ, Hastie T, Dudík M, Chee YE, Yates CJ. A statistical explanation of MaxEnt for ecologists. Diversity and Distributions. 2011; 17:43–57.

18. Rodgers WA, Panwar SH. Biogeographical classification of India. New Forest, Dehradun, India, 1988. Available from: http://wiienvis.nic.in/database/htmlpages/bioprovincemap.htm.

19. National Repository of Open Educational Resources. [cited 01 March 2019]: In NROER [Internet]. Available from: https://nroer.gov.in/home/file/readDoc/57cff54a16b51c038dedc62d/Annual%20Rainfall.svg.

20. eBird: An online database of bird distribution and abundance. eBird, Ithaca, New York [Accessed on 2 February 2019]. Database: eBird [Internet]. Available from: http://www.ebird.org.

21. Hijmans RJ, Cameron SE, Parra JL, Jones PG, Jarvis A. Very high resolution interpolated climate surfaces for global land areas. International Journal of Climatology. 2005; 25:1965–1978. Available from: www.worldclim.org

22. Bhuvan, India Geo-platform of ISRO [cited 26 February 2019]. Available from: https://bhuvan.nrsc.gov.in/bhuvan_links.php

23. USGS, Earth Resources Observation and Science (EROS) Center, SRTM Data Products [cited 2018 December 21]. In: USGS.gov [Internet]. Available from: https://www.usgs.gov/centers/eros/science/usgs-eros-archive-products-overview?qt-science_center_objects=0#qt-science_center_objects

24. Sriramadas, A. Geology of Eastern Ghats in Andhra Pradesh. Proceedings of the Indian Academy of Sciences - Section B. 1967; 66: 200–205.

25. Liu C, White M, Newell G. Selecting thresholds for the prediction of species occurrence with presence-only data. Journal of Biogeography. 2013; 40: 778–789.

26. Allouche O, Tsoar A, Kadmon R. Assessing the accuracy of species distribution models: prevalence, kappa and the true skill statistic (TSS). Journal of Applied Ecology. 2006; 43: 1223–1232.

27. Raes N, terSteege H. A null-model for significance testing of presence-only species distribution models. Ecography. 2007; 30: 727–736.

28. Sreekar R, Srinivasulu C. New site record of Yellow-throated Bulbul Pycnonotus xantholaemus from Andhra Pradesh. Indian Birds. 2010; 5: 157.

29. Indian Network for Climate Change Assessment, Ministry of Environment & Forests, Government of India. November 2010 [cited 30 April 2019]. In India environment portal [Internet]. Climate change and India: A 4×4 assessment, a sectoral and regional analysis for 2030s. Available from: http://www.indiaenvironmentportal.org.in/files/fin-rpt-incca.pdf

30. Jiguet F, Gadot AS, Julliard R, Newson SE, Couvet D. Climate envelope, life history traits and the resilience of birds facing global change. Global Change Biology. 2007; 13:1672–1684.

31. Allen, PR. Notes on the Yellow-throated Bulbul *Pycnonotus xantholaemus*. Journal of Bombay Natural History Society. 1908; 18: 905–907.

32. Bloom DE. Population dynamics in India and implications for economic growth. In: Ghate C, editor. Oxford University Press. 2011. Available from: https://core.ac.uk/download/pdf/6494801.pdf

33. Ramachandran RM, Roy PS, Chakravarthi V, Sanjay J, Joshi PK. Long-term land use and land cover changes (1920–2015) in Eastern Ghats, India: Pattern of dynamics and challenges in plant species conservation. Ecological Indicators. 2018; 85, 21–36.

34. DeFries R, Pandey D. Urbanization, the energy ladder and forest transitions in India’s emerging economy. Land Use Policy. 2010; 27: 130–138.

35. Indian Bureau of Mines. Indian Mineral Industry at a Glance 2015-16, The Mining and Mineral Statistics Division, 2018. Accessed from: https://ibm.gov.in/writereaddata/files/05092018145252IMIG-2015-16_advance%20release_mod.pdf

36. Aggregate Business International [Internet]. Booming Indian aggregates market; c2013 [cited 2019 April 20]. Available from: http://www.aggbusiness.com/sections/market-reports/features/booming-indian-aggregates-market/

37. Wilcox BA, Murphy DD. Conservation strategy: the effects of fragmentation on extinction. American Naturalist. 1985; 125: 879–887.

